# Predicting the animal hosts of coronaviruses from compositional biases of spike protein and whole genome sequences through machine learning

**DOI:** 10.1101/2020.11.02.350439

**Authors:** Liam Brierley, Anna Fowler

## Abstract

The COVID-19 pandemic has demonstrated the serious potential for novel zoonotic coronaviruses to emerge and cause major outbreaks. The immediate animal origin of the causative virus, SARS-CoV-2, remains unknown, a notoriously challenging task for emerging disease investigations. Coevolution with hosts leads to specific evolutionary signatures within viral genomes that can inform likely animal origins. We obtained a set of 650 spike protein and 511 whole genome nucleotide sequences from 225 and 187 viruses belonging to the family *Coronaviridae*, respectively. We then trained random forest models independently on genome composition biases of spike protein and whole genome sequences, including dinucleotide and codon usage biases in order to predict animal host (of nine possible categories, including human). In hold-one-out cross-validation, predictive accuracy on unseen coronaviruses consistently reached ∼73%, indicating evolutionary signal in spike proteins to be just as informative as whole genome sequences. However, different composition biases were informative in each case. Applying optimised random forest models to classify human sequences of MERS-CoV and SARS-CoV revealed evolutionary signatures consistent with their recognised intermediate hosts (camelids, carnivores), while human sequences of SARS-CoV-2 were predicted as having bat hosts (suborder Yinpterochiroptera), supporting bats as the suspected origins of the current pandemic. In addition to phylogeny, variation in genome composition can act as an informative approach to predict emerging virus traits as soon as sequences are available. More widely, this work demonstrates the potential in combining genetic resources with machine learning algorithms to address long-standing challenges in emerging infectious diseases.

## Background

The ongoing COVID-19 pandemic remains a significant public health emergency. Since the first identified cases in China in December 2019, this outbreak of respiratory disease has developed into a global crisis, with over 43 million cases worldwide to date (WHO, 2020). The causative virus was termed ‘severe acute respiratory syndrome-related coronavirus 2’ (SARS-CoV-2) (Gorbalenya et al., 2020) and is a previously unknown betacoronavirus that likely emerged through zoonotic transmission from contact with non-human animals (Andersen et al., 2020; Zhou et al., 2020). However, the precise origins of the current pandemic remain inconclusive at present (Zhang and Holmes, 2020).

Two other betacoronaviruses have zoonotically emerged to cause significant human epidemics. Severe acute respiratory syndrome-related coronavirus (SARS-CoV) emerged in China in 2002 via an intermediate host of masked palm civets (*Paguma larvata*) in live animal markets (Guan et al., 2003; Song et al., 2005), and Middle East respiratory syndrome related coronavirus (MERS-CoV) emerged in Saudi Arabia in 2012 via an intermediate host of dromedary camels (Alagaili et al., 2014; Chu et al., 2014), with considerable evidence that both originated in bats (Anthony et al., 2017; Cui et al., 2007; Hu et al., 2015; Lau et al., 2005). Four additional coronaviruses are known to be endemic within humans, causing mild common cold-like illness (*Alphacoronavirus:* Human coronaviruses 229E and NL63; *Betacoronavirus:* Human coronaviruses HKU1 and OC43).

All viruses in the family *Coronaviridae* feature similar structural proteins, including a spike glycoprotein on the outer viral surface. This protein attaches to host cell receptors and subsequently initiates cell entry. While SARS-CoV-2 shows high genetic similarity to bat coronaviruses, particularly bat coronavirus RaTG13 (matching 96% sequence identity) (Zhou et al., 2020), its spike protein instead exhibits differences among key amino acid residues of the receptor binding domain (RBD) (Andersen et al., 2020; Wan et al., 2020), the region which directly interacts with host receptors. Based on this region, SARS-CoV-2 is predicted through structural (Wan et al., 2020; Wrapp et al., 2020) and in vitro experimental models (Hoffmann et al., 2020; Letko and Munster, 2020) to have highly efficient binding to the human angiotensin-converting enzyme 2 (ACE2) receptor, a feature that has likely contributed to its efficient human-to-human transmissibility. As the key molecular determinant of host range (Graham and Baric, 2010), adaptation of coronavirus spike proteins therefore represents a key opportunity to further understand their host range constraints.

Beyond selection acting at specific loci, viral adaptation can also manifest through broad-scale genomic signatures. Viruses exhibit biased genome composition, for example, in non-uniform use of synonymous codons (Jenkins and Holmes, 2003). Furthermore, coevolution within different hosts may indirectly lead to selection for particular compositions, as reported for nucleotide and dinucleotide usage within avian and human influenzaviruses (Greenbaum et al., 2008; Rabadan et al., 2006) and codon pair usage of arboviruses within their insect vectors and mammalian hosts (Shen et al., 2015). The *Coronaviridae* are no exception - different coronaviruses (including SARS-CoV-2) vary in their genome composition, with particularly complex patterns of codon usage in spike protein coding sequences (Dilucca et al., 2020; Gu et al., 2020), which could potentially contain important evolutionary signal regarding host origin.

Machine learning has recently gained substantial attention as a methodology in comparative modelling of emerging diseases. These methods are capable of decomposing signal in high-dimensional genomic information (a limitation of regression frameworks) without the need for sequence alignment. Genomic machine learning analyses have demonstrated the ability to not only classify viruses from recurring viral genome motifs (Randhawa et al., 2020), but also classify their broad host origins (Babayan et al., 2018; Bartoszewicz et al., 2020; Young et al., 2020; Zhang et al., 2019). Specifically considering coronaviruses, support vector machines and random forests have been trained on various genomic features to predict host group, including nucleotide and dinucleotide biases (Tang et al., 2015), amino acid composition (Qiang et al., 2020) or sequence k-mers (Li and Sun, 2018). However, previous model predictions are mostly concentrated upon bats or humans, and few analyses explicitly address the spike protein (but see Li and Sun, 2018). The exact potential of genome composition to predict host origin therefore remains unclear.

We aimed to use machine learning to understand how the complex genomic signatures of coronaviruses might predict their hosts and determine the importance of such signature in the spike protein. Specifically, we trained random forest models on compositional biases for spike protein and whole genome nucleotide sequences and compared their performance. A limitation of these approaches is that model predictions can be strongly influenced by viral sampling biases and reflect virus lineage rather than host (Di Giallonardo et al., 2017). Therefore, we undersample sequences to create balanced training data and control for similarity between related sequences during hold-one-out validation. We demonstrate the use of machine learning as a reliable method to estimate host origins of future novel coronaviruses in humans and livestock.

## Methods

### Data extraction and processing

Spike protein or whole genome sequence data for coronaviruses were identified within GenBank, using search terms

‘*txid###[Organism:noexp] AND (spike[Title] OR “S gene”[Title] OR “S protein”[Title] OR “S glycoprotein”[Title] OR “S1 gene”[Title] OR “S1 protein”[Title] OR “S1 glycoprotein”[Title] OR peplomer[Title] OR peplomeric[Title] OR peplomers[Title] OR “complete genome”[Title]) NOT (patent[Title] OR vaccine OR artificial OR construct OR recombinant[Title])’*

where successive searches were conducted replacing ### with taxonomic identifiers for each species and unranked sub-species belonging to the family *Coronaviridae* within the NCBI taxonomy database (Federhen, 2012) (n = 1585 taxonomic ids total). Matching sequences were then extracted and further filtered to exclude incomplete or truncated sequences based on a) metadata labels and b) length restrictions, discarding any spike protein sequences < 2 kilobases (kb) and any whole genome sequences outside a range of 20kb – 32kb. We accepted both spike protein coding sequences within whole genome sequences and standalone complete spike protein sequences, excluding those only covering individual S1 or S2 subunits. All sequence data searching, filtering and extraction was conducted with R package ‘rentrez’ v1.2.2 (Winter, 2017; see also Brierley, 2020).

### Host classification

For each spike protein or whole genome sequence, host names were extracted from the host organism metadata field before being resolved to the standard NCBI taxonomy using the R package ‘taxizedb’ v0.1.9.93 (Chamberlain and Arendsee, 2020; see also Brierley, 2020). Host names were automatically resolved to the highest taxonomic resolution possible and any unmatched host names were resolved manually, discarding sequences with missing/unresolvable names.

We then constructed a new variable broadly describing host category of each sequence, defined at various taxonomic levels: human (species *Homo sapiens*), camelid (family *Camelidae*), swine (family *Suidae*), carnivore (order *Carnivora*), rodent (order *Rodentia*), and bird (class *Aves*). Following a previous analysis (Babayan et al., 2018), we included two categories to represent bats (order *Chiroptera*): suborder Yinpterochiroptera (families *Craseonycteridae, Hipposideridae, Megadermatidae, Pteropodidae, Rhinolophidae*, and *Rhinopomatidae*) and suborder Yangochiroptera (all other families), based on their evolutionary divergence (Tsagkogeorga et al., 2013) and differences in ecology and hosted viruses (Moratelli et al., 2015; Young and Olival, 2016). Sequences not conforming to any of the above host categories were excluded from further analysis.

## Genomic feature calculation

We then calculated several features describing genome composition biases of each spike protein and whole genome coding sequence at nucleotide, dinucleotide or codon level. Firstly, nucleotide biases were calculated as simple proportion of A, C, G or U content. Dinucleotide biases were calculated as the ratio of observed dinucleotide frequency to expected based on nucleotide frequency, following (Babayan et al., 2018):

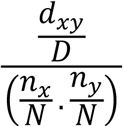

where *d*_*xy*_ denotes frequency of dinucleotide *xy, n*_*x*_ and *n*_*y*_ denote frequency of individual nucleotides *x* and *y*, and *D* denotes total dinucleotides and *N* total nucleotides for length of the given sequence. Biases were calculated separately for each dinucleotide at each position within codon reading frames (i.e. at positions 1-2, 2-3 or 3-1) as dinucleotides spanning adjacent codons may be subject to more extreme biases (Kunec and Osterrieder, 2016; Tulloch et al., 2014). Finally, Relative Synonymous Codon Usage (RSCU) was also calculated for each codon including stop codons, following (Sharp and Li, 1987):

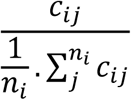

where *n*_*i*_ denotes number of codons synonymous for amino acid *i* and *c*_*ij*_ denotes frequency of *j*^*th*^ codon encoding for such amino acid. In total, this gave 116 genomic features for use in predictive models. The Effective Number of codons (ENc) (Wright, 1990) was also calculated for each sequence as a summative metric of magnitude of codon bias. All calculations for whole genomes considered nucleotide sequences as-is rather than as-read; sequence strings duplicated by frameshifting among ORF1a and ORF1b replicase protein were discarded to avoid disproportionate weighting in modelling analyses.

### Machine learning analysis

To quantify the potential for genome composition biases to predict coronavirus host category, we used random forests, an ensemble machine learning approach that aggregates over a large number of individual classification tree models (Breiman, 2001). Such predictive modelling methods are often sensitive to class imbalances in outcome variables (He and Garcia, 2009), as well as training data composition. As several viruses (e.g. Porcine epidemic diarrhea virus) and host categories (e.g. human) appeared highly over-represented, we therefore conducted stratified random undersampling prior to modelling, retaining a maximum of 20 sequences per host category per virus (Supplementary Figure S1).

Zoonotic or epizootic coronavirus sequences from their novel host (i.e. SARS-CoV, SARS-CoV-2, MERS-CoV in humans, totaling m = 9 taxonomic identifiers, full list in Supplementary Data Files; swine acute diarrhea syndrome coronavirus in swine) were held out from model training as their evolutionary signature in genome composition is much more likely to instead reflect the original or donor host, having experienced comparatively little coevolution within the novel host following cross-species transmission. We also excluded human enteric coronavirus, the zoonotic potential of which remains unclear.

All random forests were constructed using 1000 trees and implemented using R package ‘ranger’, v0.12.1 (Wright and Ziegler, 2017). Model parameters were optimized within an inner loop of 10-fold cross-validation (Supplementary Figure S1), retaining the parameter set yielding the highest prediction accuracy. This approach was repeated in an outer loop through hold-one-out validation applied to coronaviruses (Supplementary Figure S1), i.e., rather exclude a single sequence in each instance, we exclude all sequences from a single virus to control for the similarity of genome composition within-virus. This allows host predictions for novel viruses based on values of compositional bias, rather than indirectly predicting host by the proxy of viral identity.

Model performance was then assessed by applying each random forest to its respective held-out coronavirus sequences as a test set. Probabilities of host categories were obtained by dropping sequences down each individual tree model within the random forest and averaging host category prevalence of resulting terminal nodes (see Malley et al., 2012)). To investigate explanatory relationships between genomic biases and hosts of coronaviruses, variable importance (calculated as relative mean decrease in Gini impurity) and partial dependence (calculated as marginal probability of each host category) associated with each genomic feature were averaged across all random forests. Host category predictions were also generated for each zoonotic coronavirus sequence by averaging predicted probabilities across all random forests to investigate model utility in application to a newly-identified human epidemic coronavirus.

All data processing and modelling were conducted within R v4.0.0 (R Development Core Team, 2020).

## Results

### Genome composition across the *Coronaviridae*

In total, we identified n = 4960 nucleotide sequences for coronavirus spike proteins and n = 2987 whole genome sequences that met inclusion criteria. These were undersampled to n = 650 spike protein sequences from m = 225 coronaviruses and n = 511 whole genome sequences from m = 187 coronaviruses for use in further analysis (Supplementary Figure S1, Supplementary Table S1), spanning 40 identified host genera and 58 identified host species (Supplementary Data Files 1 & 2).

Broadly consistent genome composition biases were observed across the diversity of all coronavirus sequences examined. Specifically, ACU, AGA, GGU and GUU codons were over-represented among both coronavirus spike protein sequences and whole genomes (Figure 1). Hierarchical clustering based on RSCU values suggested codon usage was less distinct between genera within spike protein sequences (Figure 1A) than within whole genomes (Figure 1B). However, clear separation of deltacoronaviruses was observed for both cases, as these appeared to have less extreme biases in codon usage. This was confirmed by ENc calculation; deltacoronaviruses had higher ENc values than other coronavirus genera (Table 1). Considering dinucleotide biases, compositional bias was typically more extreme for dinucleotides spanning adjacent codons, i.e. position 3-1 (Supplementary Figure S2), and the characteristic coronavirus CpG suppression was also observed.

**Table 1.** ENc values across coronavirus genera. Effective Number of Codons (ENc) for coronaviruses, stratified by genus and genome sequence type. ENc values are calculated as grand means, i.e. mean ENc was calculated per coronavirus by averaging sequences before means of means were calculated per genus by averaging coronaviruses. SD denotes standard deviation.

|  | Spike protein sequences |  | Whole genome sequences |  |
| --- | --- | --- | --- | --- |
| Genus | Mean | SD | Mean | SD |
| *Alphacoronavirus* | 48.65 | 3.12 | 45.02 | 3.00 |
| *Betacoronavirus* | 47.77 | 5.04 | 47.03 | 4.79 |
| *Gammacoronavirus* | 46.36 | 1.83 | 46.11 | 0.74 |
| *Deltacoronavirus* | 54.79 | 3.43 | 51.75 | 3.22 |
| (unclassified) | 47.89 | 3.41 | 48.56 | 3.04 |
| **total** | 48.77 | 4.36 | 46.81 | 4.22 |

**Figure 1.**
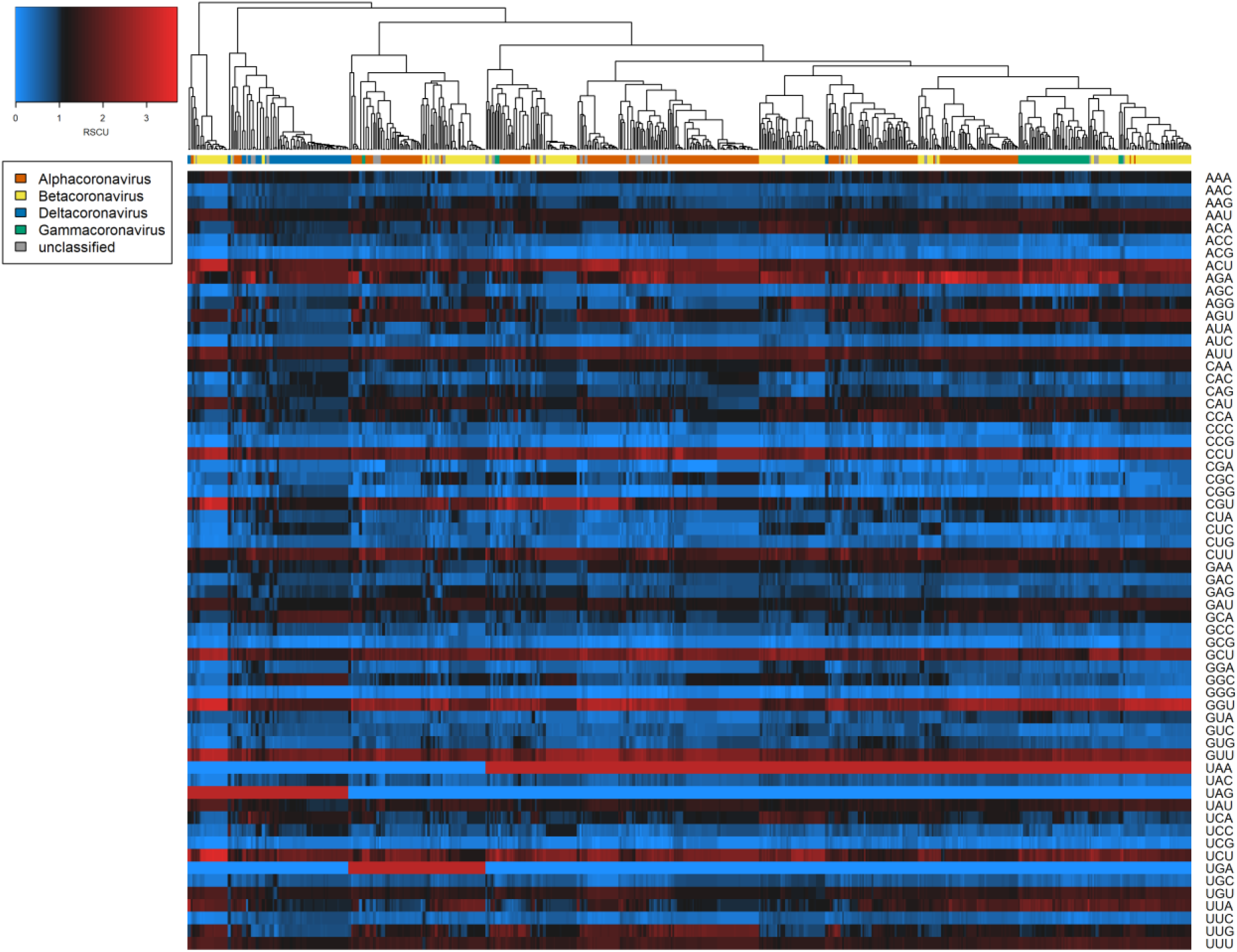
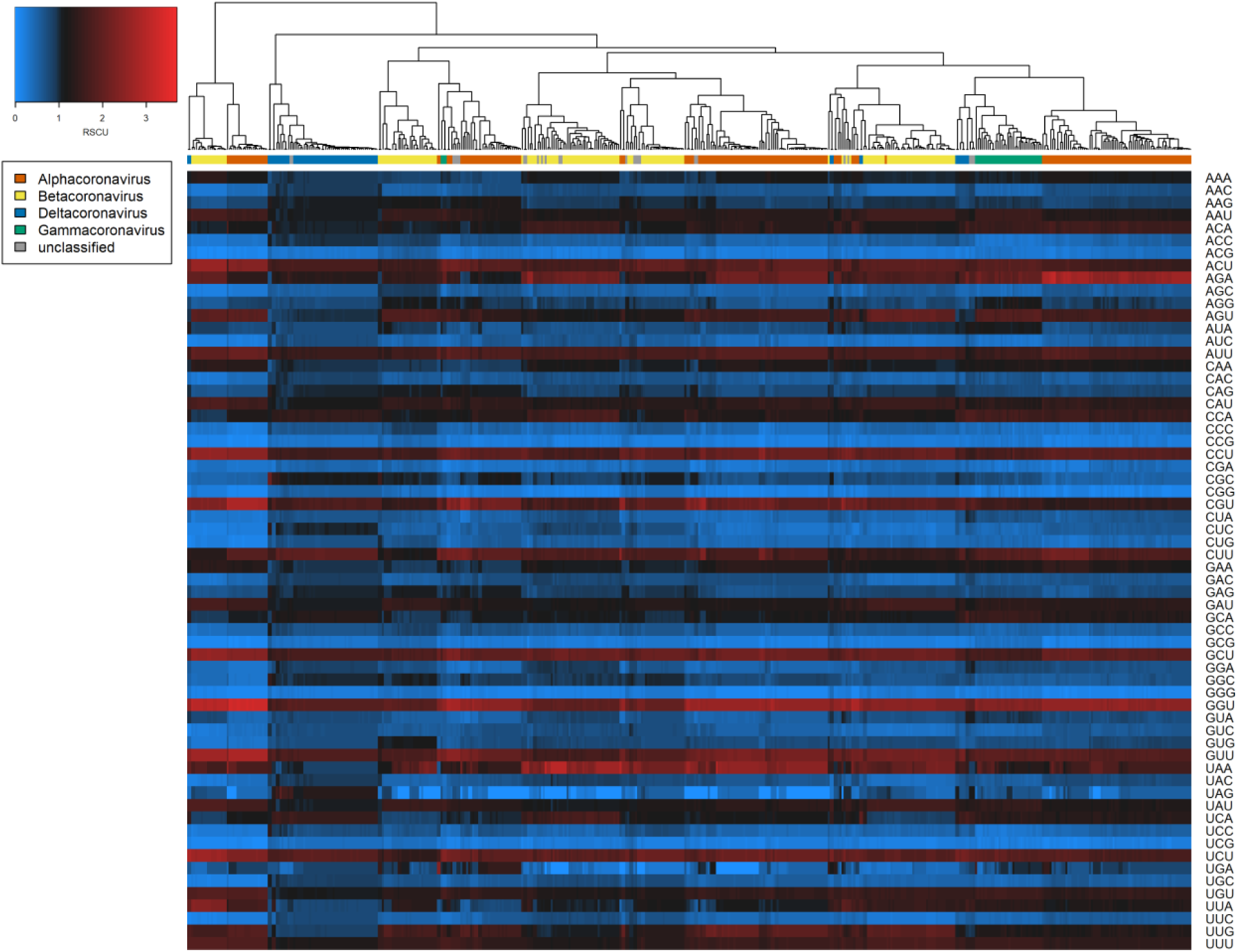
Codon biases (RSCU) across coronavirus genome sequences examined. Heatmaps of coronavirus codon usage bias (RSCU) associated with each codon in each A) spike protein sequence (n = 650) and B) whole genome sequence (n = 511). Main colour scale denotes RSCU value, a null value of 1 (black) indicating no difference in codon usage from expectation, with blue and red representing under-or over-representation respectively. Dendrogram colour bar denotes taxonomic genus.

### Host predictions of random forest models

Random forest models trained on nucleotide, dinucleotide and codon bias features of spike protein sequences predicted coronavirus hosts with 73% accuracy during hold-one-out cross-validation (Table 2). Genome composition of spike proteins appeared just as informative as whole genomes despite being much smaller in sequence length, as both models achieved very similar performance in all diagnostic measures (Table 2).

**Table 2.** Predictive performance of random forest models. Model diagnostics describing overall performance when applied to predict host category of held-out coronaviruses. CI denotes confidence interval, Kappa denotes Cohen’s Kappa statistic, mAUC denotes multiclass area-under-curve statistic. F1micro and F1macro denote F1 scores calculated using micro-averaging (performance on each host category weighted proportionally) and macro-averaging (performance on each host category weighted equally), respectively.

| Predictor features | Accuracy (95% CI) | Kappa | mAUC | F1<sub>micro</sub> | F1<sub>macro</sub> |
| --- | --- | --- | --- | --- | --- |
| Spike protein | 0.735 (0.700, 0.769) | 0.696 | 0.898 | 0.848 | 0.757 |
| Whole genome | 0.728 (0.687, 0.766) | 0.688 | 0.902 | 0.843 | 0.758 |

Patterns of host-specific predictive performance were evident during hold-one-out cross-validation. Random forests trained on both spike protein and whole genome sequence compositional features most easily distinguished bird, carnivore and rodent host categories (Figure 2, see also Supplementary Tables S2 & S3). Less powerful predictive performance was obtained for livestock (i.e., swine and camelid) host categories with these sequences often predicted as having bat (suborder Yangochiroptera) hosts, including all MERS-CoV sequences sampled from camels.

**Figure 2.**
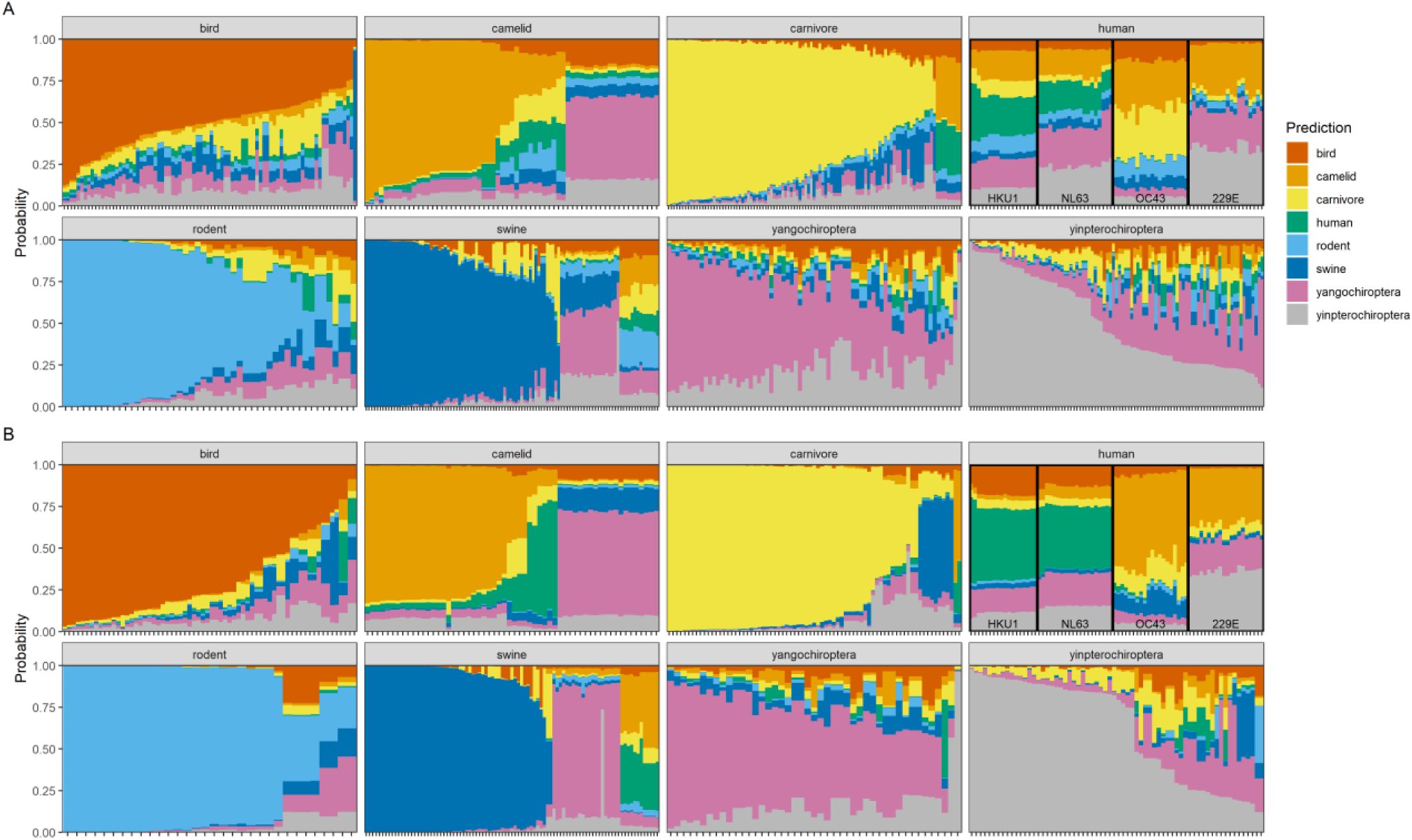
Random forest host predictions based on coronavirus genome composition. Stacked bar plots of predicted probabilities of each host category for coronavirus sequences. Predictions were obtained from ensemble random forest models trained on A) spike protein and B) whole genome composition features. Panels depict sequences from each metadata-derived host category and colour coding denotes model-predicted host category. Stacks represent individual coronavirus sequences, ordered from largest to smallest probability of the correct host, i.e. greater panel area matching the correct host category indicates better overall model performance. Non-zoonotic coronavirus sequences originating from humans (human coronaviruses HKU1, NL63, OC43, 229E) are labelled for clarity.

Human host origins appeared particularly difficult to characterise, with model-predicted hosts appearing more uncertain using spike protein features than whole genome features (Figure 2); while human coronaviruses HKU1 and NL63 were more confidently correctly classified based on whole genomes, human coronaviruses OC43 and 229E were more confidently misclassified as having camelid or Yinpterochiroptera hosts. The reciprocal also only occurred using whole genomes, i.e. several camel coronaviruses were predicted to have human hosts.

We then applied random forests to those sequences of zoonotic viruses sampled from humans and excluded from model training: SARS-CoV, SARS-CoV-2, and MERS-CoV (Supplementary Figure S1, Supplementary Data Files 3 & 4). As these viruses have experienced little coevolution following zoonotic spillover, their genome composition signal likely gives indications about their ultimate or proximate animal host origins. MERS-CoV was overwhelmingly predicted to have camelid hosts (Figure 3) and SARS-CoV was predicted with less certainty as having carnivore hosts, consistent with the respective known intermediate hosts of camels and palm civets (order *Carnivora*). Contrastingly, SARS-CoV-2 was predicted mostly strongly to have a bat (suborder Yinpterochiroptera) host. Host predictions for zoonotic viruses were consistent between models using spike protein and whole genome features (Figure 3).

**Figure 3.**
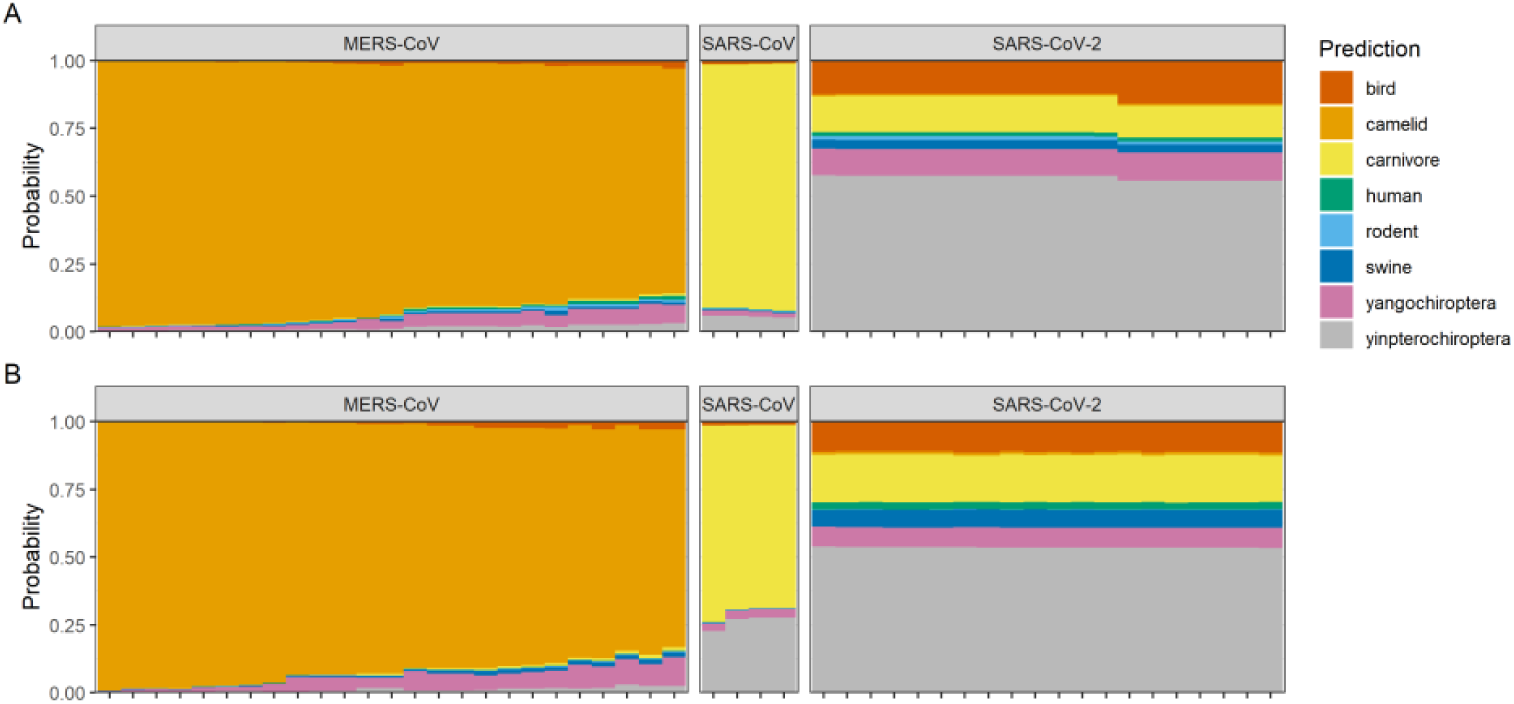
Random forest predictions based on zoonotic coronavirus genome composition. Stacked bar plots of predicted probabilities of each host category for zoonotic coronavirus sequences sampled from humans. Predictions were obtained from ensemble random forest models trained on A) spike protein and B) whole genome composition features. Colour coding denotes model-predicted host category. Stacks represent individual coronavirus sequences.

### Variable importance of random forest models

The most informative genomic features towards predicting coronavirus hosts were a mixture of dinucleotide and codon biases (Figure 4), with dinucleotide biases appearing slightly more informative for spike protein sequences and codon biases appearing slightly more informative for whole genome sequence. However, predictive power of individual genomic features did not hold between spike protein and whole genome sequences; only weak correlation was observed between ranked variable importance from both analyses (Spearman’s rank, ρ = 0.191, p = 0.042; Figure 4, see also Supplementary Figures S4 & S5). Partial dependence plots suggested the strongest individual discriminating feature to be GG dinucleotides at positions 1-2; an overrepresentation of this dinucleotide within the spike protein sequence clearly distinguished bird hosts from mammalian hosts (Supplementary Figure S4), consistent with the greatest predictive performance for bird coronaviruses (Supplementary Table S2).

**Figure 4.**
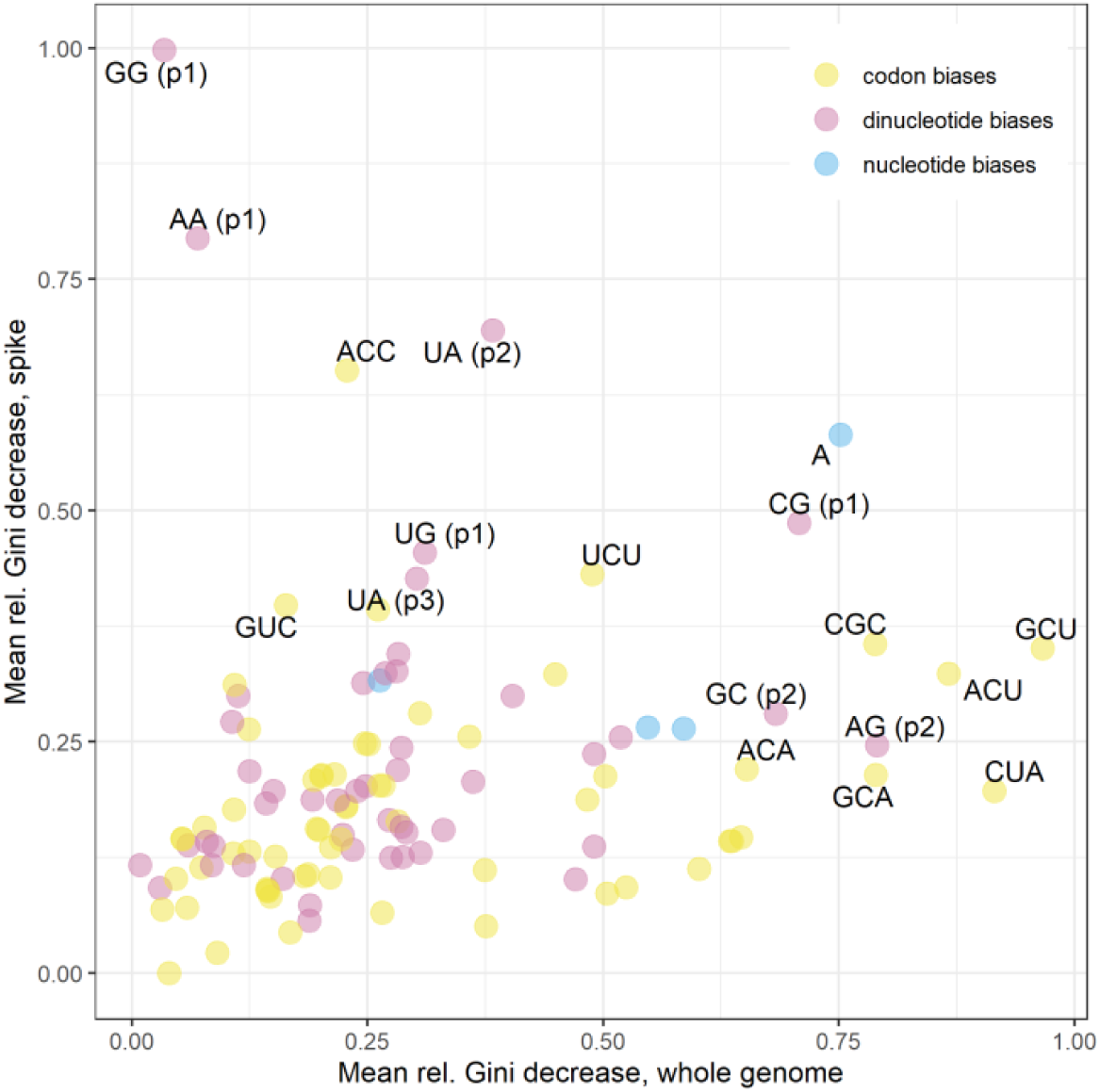
Variable importance of genomic features. Variable importance of genome composition features in ensemble random forest models predicting coronavirus host category from whole genome sequences (x axis) and spike protein sequences (y axis), with labelling of top ten most informative features from both analyses. Points denote mean values of relative decrease in Gini impurity associated with each feature across A) m = 225 and B) m = 187 random forests during hold-one-out validation. Colour key denotes genomic feature type.

## Discussion

We observe biased dinucleotide and nucleotide usage across the family *Coronaviridae*, and demonstrate that these genome composition biases contain sufficient evolutionary signal such that they can predict animal host origin. We show that training random forests on these features of spike proteins is equally as informative as using whole genome sequences in predicting hosts of novel (i.e., unseen) coronaviruses, with bird, carnivore and rodent viruses having the highest prediction accuracy. When applied to human coronavirus sequences from previous epidemics (SARS-CoV, MERS-CoV), random forest model predictions consistently represented the intermediate hosts. In the case of SARS-CoV-2, where the exact transmission pathway remains unknown, models predicted sequences to have a bat host (suborder Yinpterochiroptera).

### Variability in genome composition

Among our dataset of 225 coronaviruses, we observed A- and U-ending codons to be overrepresented and G- and C-ending codons to be underrepresented (Figure 1), a commonly noted trait in other studies (Dilucca et al., 2020; Tort et al., 2020). Elsewhere, CpG dinucleotide bias has been proposed as a specific determinant of host (and tissue) range of coronaviruses (Xia, 2020), on the basis that CpG dinucleotides are targeted by zinc finger antiviral proteins and their suppression is therefore linked to immune evasion. We observed consistent CpG suppression (Supplementary Figure S2) and CG dinucleotides (positions 1-2) ranked 6^th^ and 8^th^ in feature importance for spike proteins and whole genomes (Figure 4), respectively.

However, spike proteins display different patterns of codon usage from other viral proteins (Gu et al., 2020), reflected in the lack of correlation between genome composition feature importance in random forests trained on spike proteins versus whole genomes (Figure 3, see also Supplementary Figures S4 & S5). This indicates spike proteins contain complex and distinct evolutionary signatures indicative of host-virus coevolution, supporting our approach in using many features beyond single dinucleotides (Pollock et al., 2020).

### Model predictions of human coronaviruses

During model validation, human hosts appeared more challenging to correctly predict than other host types. Although the endemic human coronaviruses are common, they are also thought to have their ultimate evolutionary origins within non-human animals (Cui et al., 2007), which may explain this difficulty. In particular, human coronaviruses NL63 and HKU1 were more consistently predicted as having human hosts than human coronaviruses OC43 and 229E, especially when using whole genome sequences (Figure 2). Human coronavirus NL63 is estimated to have a more ancient common ancestor with bat coronaviruses than 229E (Huynh et al., 2012; Pfefferle et al., 2009), implying longer coevolutionary history within human hosts has resulted in a more consistently identifiable genomic signature.

Although several mutations of SARS-CoV-2 are becoming fixed in the population, e.g. D614G in the spike protein (Grubaugh et al., 2020), the virus has experienced only weak purifying selection (MacLean et al., 2020) and sequences remain extremely similar over the course of the pandemic. As such, our approach cannot identify host adaptation “in real-time”; rather, we examine variation generated over much longer macroevolutionary histories.

Instead, we would expect viruses that have transmitted cross-species more recently to retain the genome composition signature of their previous hosts, having experienced little coevolution within the novel host. Applying our finalised models to zoonotic human virus sequences may therefore give an indication of their proximate or ultimate animal host origin (Figure 3).

Human sequences of SARS-CoV were predicted to have a carnivore host, consistent with the known intermediate host of palm civets (*Paguma larvata)*. Much previous work has shown human and civet SARS-CoV sequences to have high similarity, with adaptive mutations concentrated within the spike protein (specifically, the receptor binding domain) (Graham and Baric, 2010; Guan et al., 2003; Song et al., 2005), which may explain the stronger prediction of carnivore hosts when using spike protein sequences. Similarly, human sequences of MERS-CoV were strongly predicted as having camelid hosts, consistent with the intermediate host of dromedary camels. The detection of evolutionary signatures corresponding to these intermediate hosts implies that these coronaviruses circulated in those hosts for sufficient time for coevolution to shape genome composition before cross-species transmission to humans. For MERS-CoV, camel infections have been recognised as far back as at least the 1980s (Müller et al., 2014; Sabir et al., 2016).

The origins of SARS-CoV-2 have been heavily speculated upon since its discovery, though there remains no compelling evidence towards the animal source of the first human infections. Our random forest models trained on genome composition of both spike protein and whole genome sequences predicted SARS-CoV-2 as having a bat host (suborder Yinpterochiroptera). Alignment-based and phylogenetic approaches showed the most closely related virus to be bat coronavirus RaTG13, a virus sampled from a horseshoe bat (*Rhinolophus affinis*) belonging to this suborder (Zhou et al., 2020), and more widely, the *Rhinolophidae* family are the most likely ancestral hosts of the *Sarbecovirus* genus (Latinne et al., 2020).

While our model predictions support bats as the ultimate origin of SARS-CoV-2, the involvement of intermediate hosts remains unclear. Although the Malayan pangolin (*Manis javanica*) was proposed early in the pandemic (Liu et al., 2020; Xiao et al., 2020), recent analyses have argued there is absence of evidence for this (Boni et al., 2020; Zhan et al., 2020). The methods used here are unable to identify intermediate hosts without sufficient sequence availability, and lack of such from pangolins (order *Pholidota*) preclude us from directly testing this hypothesis. However, selection analyses indicate SARS-CoV-2 could reasonably have exhibited efficient human infectivity and human-to-human transmissibility following direct transmission from bats (MacLean et al., 2020; Zhan et al., 2020), i.e., without strict need for prolonged selection within an intermediate host.

### Future directions

A natural comparison to these methods is phylogenetic analyses, which can estimate traits such as host type from reconstructing viral ancestry based on sequence similarity. There is challenging confounding between molecular characteristics and sequence similarity, i.e., variation in genome composition may actually be predictive of viral lineage rather than host type (Di Giallonardo et al., 2017), essentially acting as a proxy for phylogenetic relatedness. To separate these signals, phylogeny needs to be considered in model construction (Young et al., 2020); by using a cross-validation procedure holding out individual coronaviruses rather than individual sequences, we attempt to distinguish genomic signatures arising from convergent evolution within specific hosts, rather than from viral similarity. A more generalised scope of study across multiple viral families would allow holdout of entire families during cross-validation (Young et al., 2020), further removing any phylogenetic proxy effects.

Additional challenges are created by the unavoidable, systematic gaps in sampling coverage. For example, disproportionate sampling to identify viruses in wildlife similar to those already known to affect humans or livestock may introduce bias to predictive models. Although we address this by undersampling, our model predictions, particularly for zoonotic coronaviruses, are likely influenced by the range of known viruses with available sequence data. These issues highlight the need for a careful choice of training dataset in modelling studies, but also for wider sampling and surveillance of coronaviruses among the wild virome (Carroll et al., 2018), especially considering their high public health risks.

Although we focus on compositional features, predictive approaches using other sequence properties may improve more mechanistic understanding of host range. For example, amino acid composition and physiochemical similarity between contiguous amino acid residues can predict human origin of coronavirus spike sequences (Qiang et al., 2020). Similarly, Young et al. (2020) have recently demonstrated the use of multiple types of genomic features in combination to predict infected hosts and found physicochemical classification of amino acid k-mers to achieve similar predictive power to nucleotide k-mers. More widely, hydrophobic and hydrophilic composition of host receptors shows some predictive signal towards virus sharing (Bae and Son, 2011), hydropathy being of mechanistic importance during virus-receptor binding, e.g. for murine coronavirus (Thackray et al., 2005). These properties could be used as additional features and improve machine learning model-derived predictions.

This emphasises an additional key question for future modelling studies distinct from host origin - whether genomic traits can predict the zoonotic potential of newly discovered animal coronaviruses. As this is strongly determined by molecular mechanisms of virus-host receptor interactions (Graham and Baric, 2010), these predictions may be best inferred by model frameworks combining genomic features of both spike proteins and host receptors.

## Conclusion

By training machine learning models on genome composition across the *Coronaviridae*, we demonstrate a detectable evolutionary signature predictive of host type rooted in a region of the genome that is key to host shifts. Our random forest predictions add to the growing evidence COVID-19 ultimately originated within bats, though further work is needed to understand the potential for intermediate hosts. Characterising spike proteins (and by extension, their interaction with host receptors) may provide a fruitful path to further understanding zoonosis risk among coronaviruses.

## Acknowledgements

We thank Cillian Courtney, Maya Wardeh, and Matthew Baylis for helpful support and commentary on this work. LB acknowledges funding from a Medical Research Council Skills Development Fellowship award, grant number MR/T027355/1.

