## Supplementary Material (Data Captions, Tables, Figures) for "Predicting the animal hosts of coronaviruses from compositional biases of spike protein and whole genome sequences through machine learning"

#### Supplementary Data

**Supplementary Data Files 1 & 2. Spike protein and whole genome sequences used in analysis with metadata and model-predicted host category.**

List of non-zoonotic coronavirus genome sequences used in spike protein (Data File 1) and whole genome machine learning analysis (Data File 2) along with genus, GenBank accession ID, metadata-derived host category and predicted host category from random forest when respective coronavirus was held-out during validation. Coronavirus names refer to taxonomic entities as defined by NCBI taxonomy (whether species or unranked sub-species). Model predictions are given as single category with highest probability, along with probabilities associated with each individual category.

**Supplementary Data Files 3 & 4. Spike protein and whole genome sequences of zoonotic human coronaviruses with metadata and model-predicted host category.**

List of zoonotic coronavirus genome sequences sampled from humans used in spike protein (Data File 3) and whole genome machine learning prediction (Data File 4) along with genus, GenBank accession ID, and predicted host category, aggregating over all random forests. Coronavirus names refer to taxonomic entities as defined by NCBI taxonomy (whether species or unranked sub-species). Model predictions are given as probabilities associated with each individual category.

#### Supplementary Tables

**Supplementary Table S1. Number of coronavirus sequences represented per host category.**
Number of genome sequences and unique coronavirus taxonomic ids sourced from each host category within dataset of coronavirus spike proteins and whole genome sequences. Note that although most coronaviruses were only known to infect a single host category, several coronaviruses infected multiple host categories and are represented multiple times among unique taxonomic ids.

|  | **Spike protein dataset** | | **Whole genome sequence dataset** | |
| --- | --- | --- | --- | --- |
| **Host category** | **No. sequences** | **No. coronaviruses** | **No. sequences** | **No. coronaviruses** |
| bird | 84 | 24 | 66 | 23 |
| camelid | 63 | 7 | 58 | 7 |
| carnivore | 103 | 48 | 75 | 41 |
| human | 78 | 4 | 78 | 4 |
| rodent | 49 | 22 | 32 | 18 |
| swine | 104 | 23 | 91 | 22 |
| yangochiroptera | 72 | 49 | 45 | 33 |
| yinpterochiroptera | 97 | 48 | 66 | 39 |
| **total** | 650 | 225 | 511 | 187 |

**Supplementary Table S2. Predictive performance of random forest models using spike protein predictor features, stratified by host category.**
Model diagnostics describing overall performance when applied to predict host category of held-out coronaviruses. Balanced accuracy denotes 0.5*(sensitivity + specificity).

| **Host category** | **Balanced accuracy** | **Precision** | **Recall** | **F1 score** |
| --- | --- | --- | --- | --- |
| bird | 0.976 | 1.000 | 0.952 | 0.976 |
| camelid | 0.796 | 0.615 | 0.635 | 0.625 |
| carnivore | 0.932 | 0.852 | 0.893 | 0.872 |
| human | 0.613 | 0.857 | 0.231 | 0.364 |
| rodent | 0.951 | 0.818 | 0.918 | 0.865 |
| swine | 0.824 | 0.958 | 0.654 | 0.777 |
| yangochiroptera | 0.881 | 0.445 | 0.903 | 0.596 |
| yinpterochiroptera | 0.830 | 0.673 | 0.722 | 0.697 |

**Supplementary Table S3. Predictive performance of random forest models using whole genome predictor features, stratified by host category.**
Model diagnostics describing overall performance when applied to predict host category of held-out coronaviruses. Balanced accuracy denotes 0.5*(sensitivity + specificity).

| **Host category** | **Balanced accuracy** | **Precision** | **Recall** | **F1 score** |
| --- | --- | --- | --- | --- |
| bird | 0.955 | 1.000 | 0.909 | 0.952 |
| camelid | 0.734 | 0.457 | 0.552 | 0.500 |
| carnivore | 0.918 | 0.969 | 0.840 | 0.900 |
| human | 0.730 | 0.760 | 0.487 | 0.594 |
| rodent | 0.935 | 0.933 | 0.875 | 0.903 |
| swine | 0.790 | 0.789 | 0.615 | 0.691 |
| yangochiroptera | 0.910 | 0.442 | 0.933 | 0.600 |
| yinpterochiroptera | 0.882 | 0.757 | 0.803 | 0.779 |

### Supplementary Figures

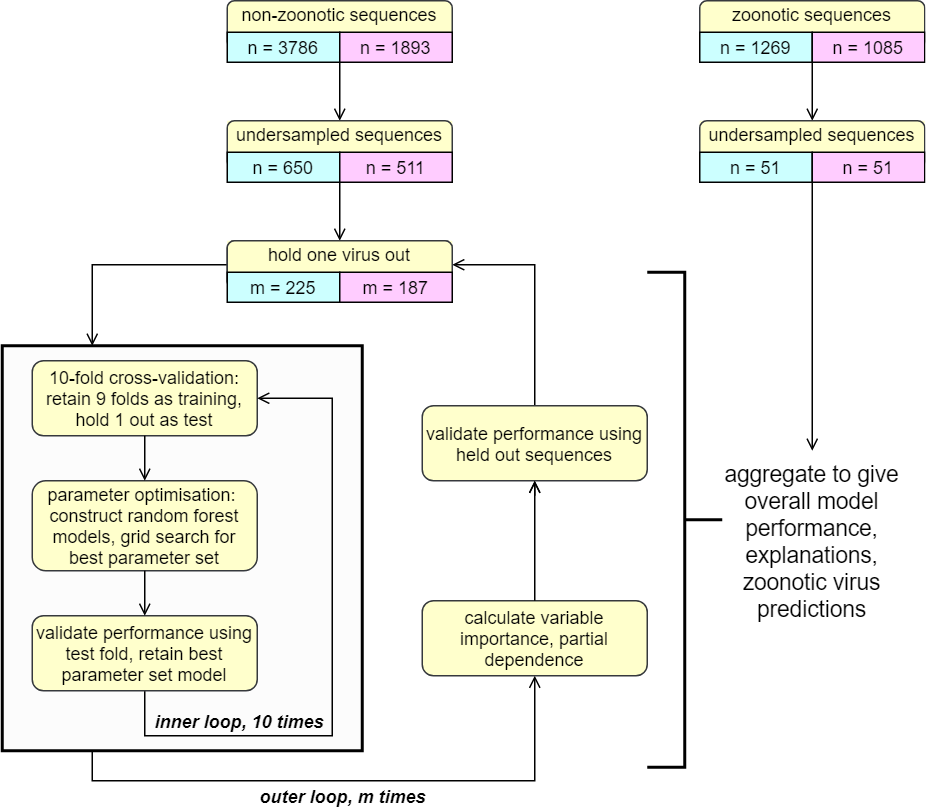

**Supplementary Figure S1. Structured data handling and machine learning modelling process.**
Flowchart indicating process of data handling, undersampling and machine learning modelling, distinguishing the inner loop (10-fold cross-validation) with the aim of optimising model parameters from the outer loop (hold-one-out cross-validation applied at coronavirus level) with the aim of validating model performance. Non-zoonotic coronavirus sequences used for model training are also distinguished from zoonotic coronavirus sequences sample from humans used for model prediction only. Blue and pink boxes denote spike protein sequence dataset and whole genome sequence dataset respectively, while n and m indicate number of individual sequences and individual coronaviruses (either species or unranked subspecies determined by taxonomic id) respectively.

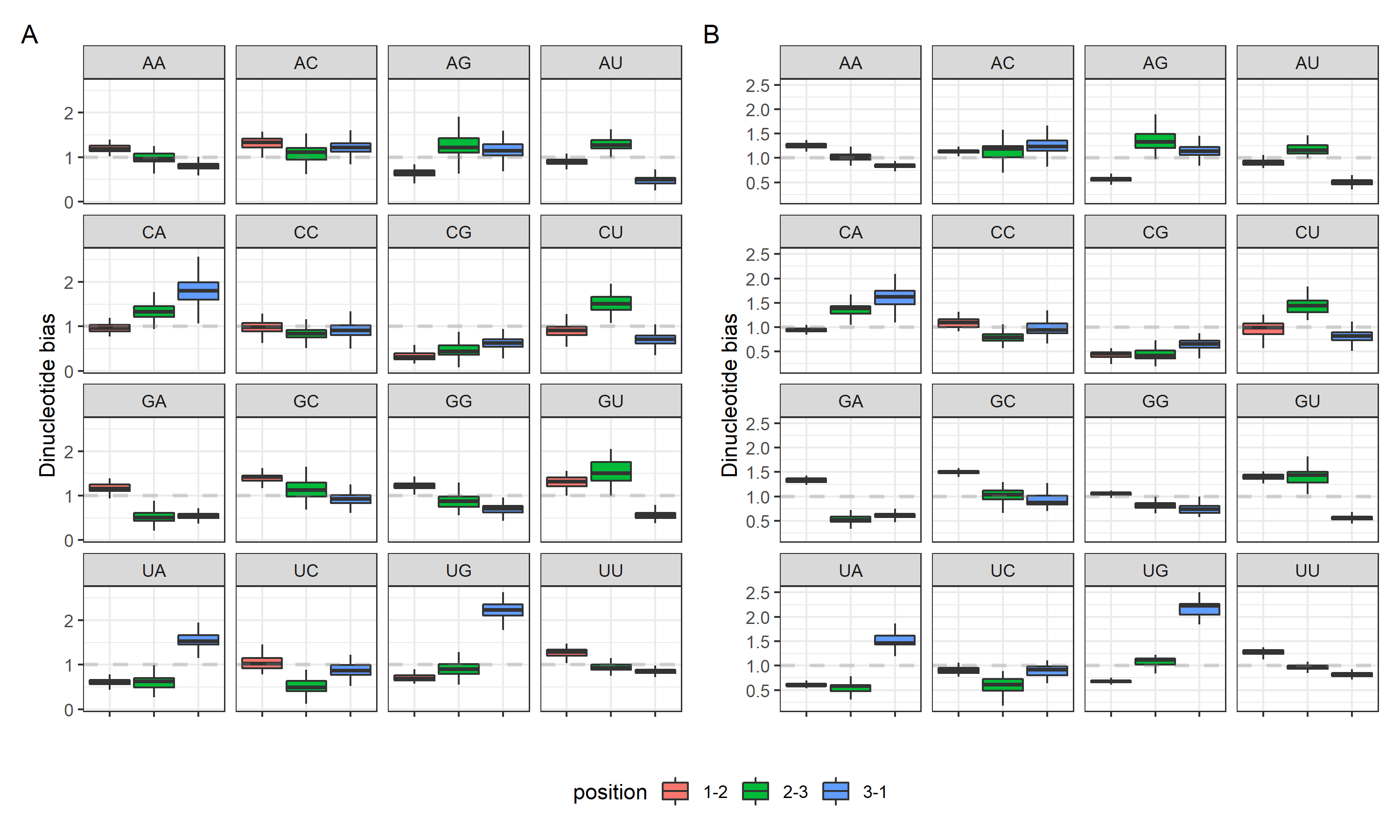

**Supplementary Figure S2. Dinucleotide biases by position within codon reading frames.**
Calculated dinucleotide biases for coronavirus A) spike protein and B) whole genome sequences. Boxes denote median and upper/lower quartiles, with whiskers extending to 1.5*IQR. Colour denotes position of dinucleotide within codon reading frames, i.e. blue boxes denote dinucleotides spanning adjacent codons (position 3-1). Dashed grey line denotes null value of 1, indicating no difference in dinucleotide usage from expectation.

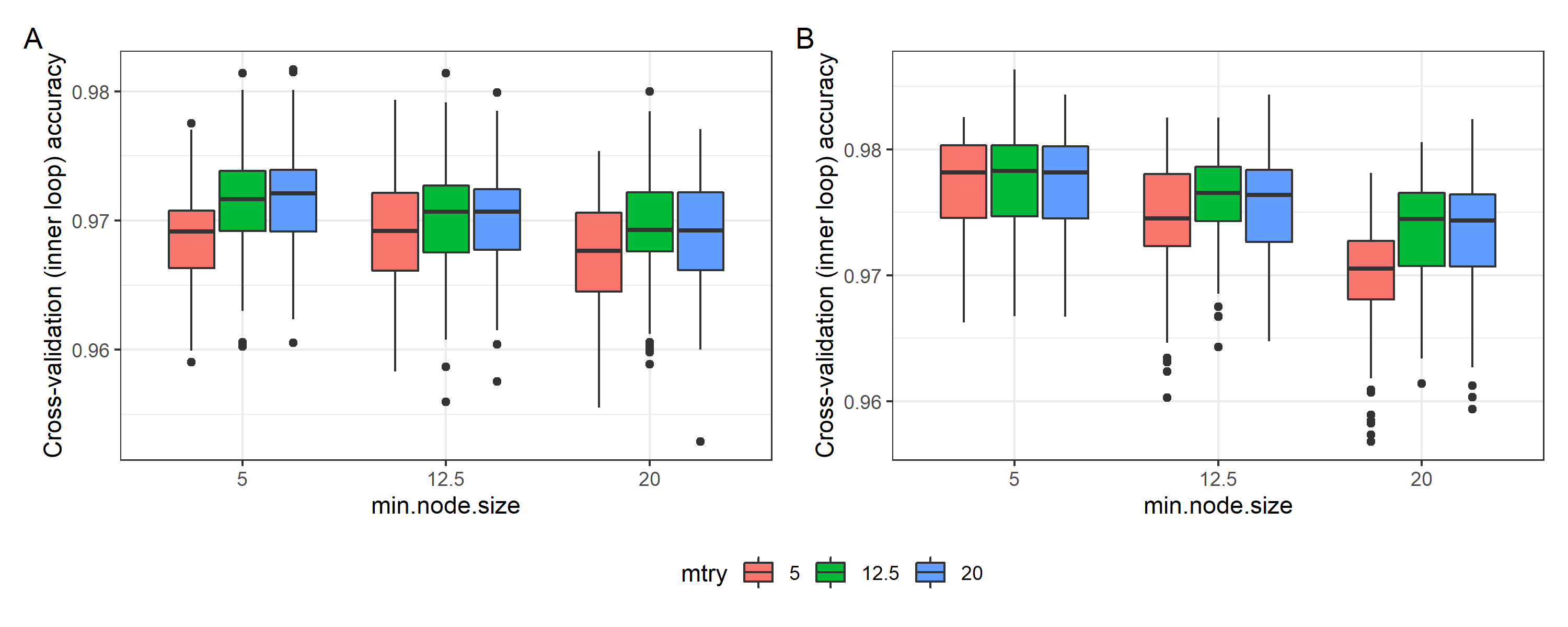

**Supplementary Figure S3. Performance of random forests during parameter optimisation.**
Performance of random forest models during 10-fold cross-validation for parameter optimisation. Y axis denotes prediction accuracy on test fold, X axis denotes minimum number of genome sequences in nodes at which tree algorithm continues to split, and colour denotes number of genomic features randomly considered at each splitting point. Boxes denote median and upper/lower quartiles, with whiskers extending to 1.5*IQR.

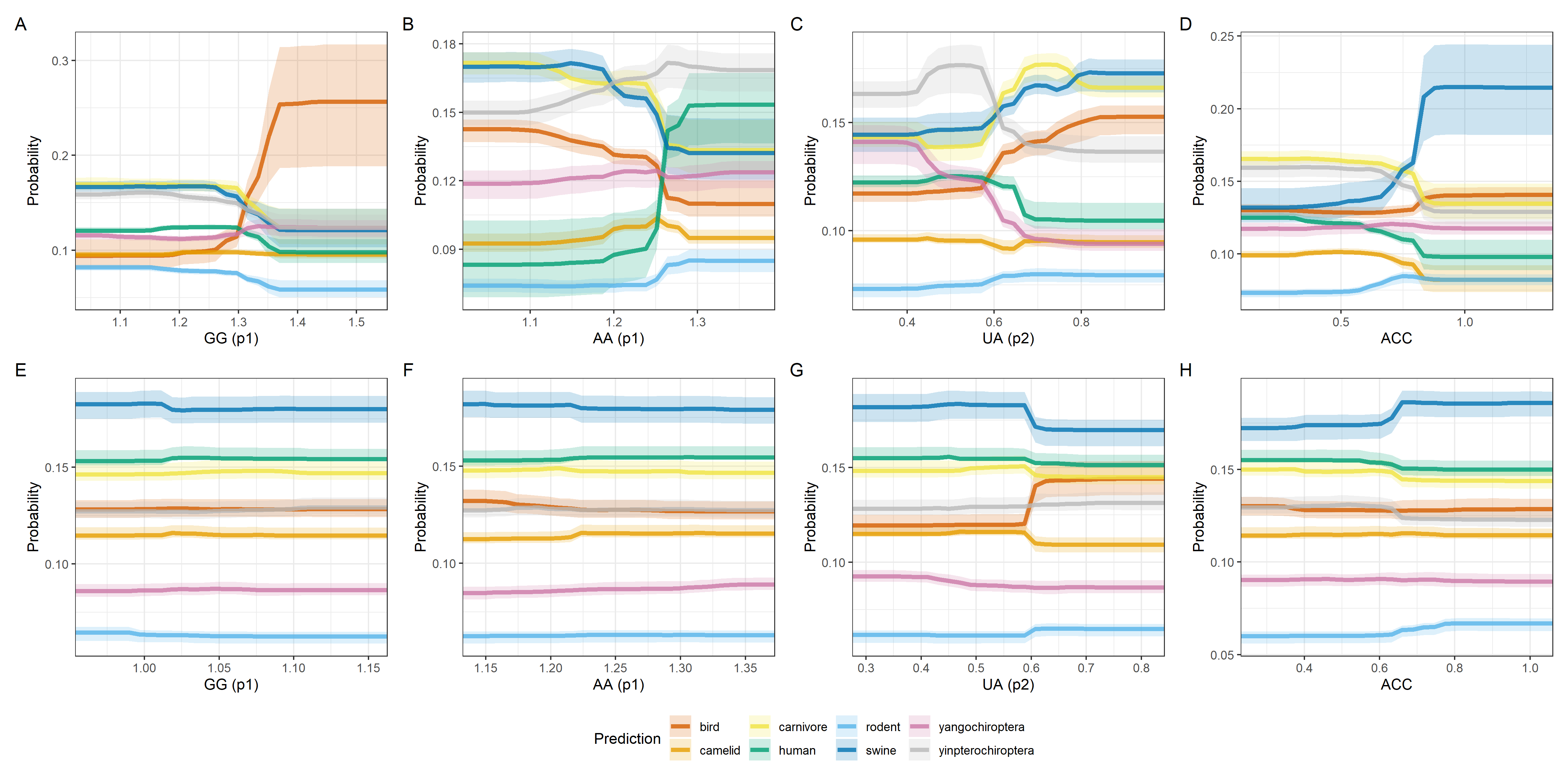

**Supplementary Figure S4. Partial dependence plots for most informative genomic features of spike proteins.**
Model-predicted marginal probability of each coronavirus host category as functions of the four most informative genome composition bias features of spike protein sequences. A) – D) depict random forests trained on spike protein sequences and E) – H) depict probabilities as functions of the same features within random forests trained on whole genome sequences for comparison. Lines denote median values across A) – D) m = 225 and E) – H) m = 187 random forests during hold-one-out cross-validation. Shaded areas denote 2.5^th^ and 97.5^th^ percentiles. Colour key denotes host category.

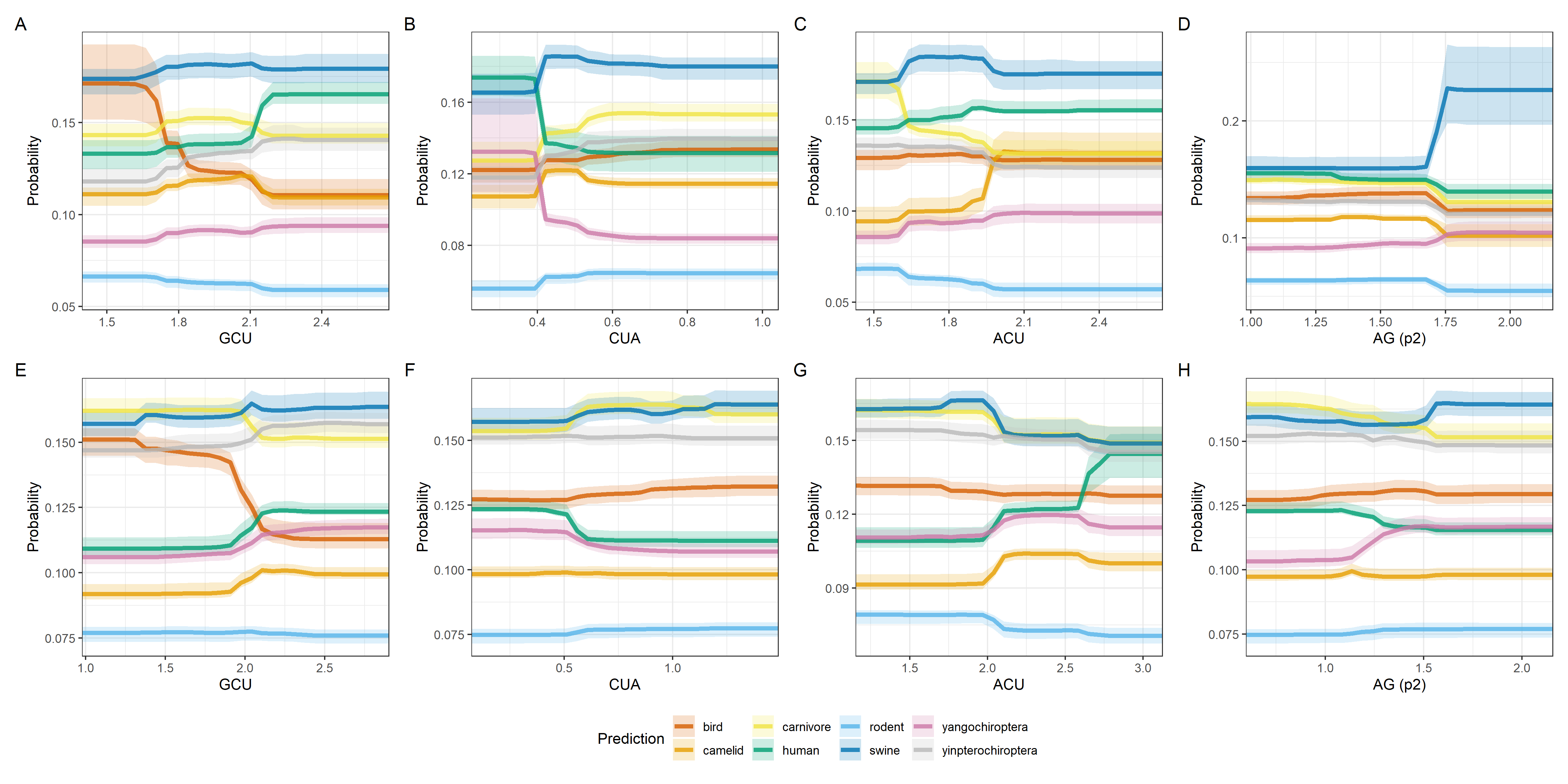

**Supplementary Figure S5. Partial dependence plots for most informative genomic features of whole genome sequences.**
Model-predicted marginal probability of each coronavirus host category as functions of the four most informative genome composition bias features of whole genome sequences. A) – D) depict random forests trained on whole genome sequences and E) – H) depict probabilities as functions of the same features within random forests trained on spike protein sequences for comparison. Lines denote median values across A) – D) m = 187 and E) – H) m = 225 random forests during hold-one-out cross-validation. Shaded areas denote 2.5^th^ and 97.5^th^ percentiles. Colour key denotes host category.
